# Long-term associations and insights on social structure of the Humpback whales in Prince William Sound, Alaska

**DOI:** 10.1101/2020.03.02.972828

**Authors:** Olga von Ziegesar, Shelley Gill, Beth Goodwin

**Affiliations:** Winged Whale Research, Homer, Alaska, United States

## Abstract

Humpback whale (*Megaptera novaeangliae*) social structure is more complex than previously thought. Because of the fluid “fission-fusion” nature of their relationships: individuals foraging, traveling, and socializing with a number of animals, where associations form and are broken numerous times, little has been confirmed about their long-term associations. Humpback whales of the North Pacific Ocean migrate annually between tropical breeding areas to northern latitudes where they congregate and feed. The purpose of this study was to explore the social and feeding habits of the summer population of humpback whales returning to Prince William Sound (PWS) in the south central coast of Alaska. Fluke photographs of pigmentation patterns were used to document individual whales between the years 1983 and 2009 to determine, population characteristics, reproductive rates, long-term associations, feeding habits and spatial partitioning. During the 27 year study period there were 3,017 encounters with 405 unique whales. Forty of these whales (9.88%) had long sighting histories, showing strong site fidelity. Association indices for all pairs of whales were calculated. Long-lasting associations were found between thirty-two of the forty whales. Two distinct groups were determined by the highest association coefficients. Although the overall ranges of the two groups overlapped, they did not often mingle and offspring did not join their maternal group. All but two females had enduring bonds with at least one male. Associate males were sometimes found at a distance from others of their “clan” and would rejoin periodically. Two whales from one of these clans were found together in Hawaiian waters, a male escorting a female with a newborn calf, suggesting these long lasting associations endure through migration and into the southern breeding areas. Optimal observation conditions of a small population of humpback whales in sheltered waters allowed the discovery of two social groups enduring almost three decades.

## Introduction

Humpback whales (*Megaptera novaeangliae*) have often been described as having fission-fusion relationships: pairs coming together and splitting again, with polygynous mating habits and no long-lasting associations [1–7]. Many earlier studies of humpback whales concluded that the social organization in humpback whales is restricted to unstable groups with few exceptions [8–11]. This twenty-seven consecutive year study in the sheltered waters of Prince William Sound, Alaska (PWS) demonstrated that long-term associations in two small groups of humpback whales existed.

Lasting social groups are documented in many large mammal species from terrestrials such as elephants, wolves and primates to marine cetaceans such as killer whales, sperm whales and dolphins. However, little is understood about the social order of the Mysticete (baleen) whales including the North Pacific humpback whales. While humpback whales’ social framework may not be as structured as the tight family units of the Odontocete (toothed) whales, this study shows they do have organized social groups.

Most Pacific humpback whales return habitually to specific summer feeding areas at northern latitudes after a long migration from tropical winter breeding grounds [12]. This study took place in Prince William Sound (PWS) in the northern apex of the Gulf of Alaska (Fig. 1) from May to October 1983 to 2009. The Sound is comprised of many sheltered bays and passages enabling an opportunity to observe whale habits in calm water. Photographs of the unique pigmentation patterns on the underside of each whale’s tail [13] were catalogued [14] and used to identify and count individual animals and to document their social behavior. About 100 individual whales visited PWS annually, many of which were returning whales. [15,16]. During the study period 405 individual whales were documented of which only 40 had long sighting histories. Two groups or “clans” of whales became apparent. These whales usually fed in different parts of PWS and rarely intermingled. Long-lasting relationships between male and female whales existed within both clans indicating humpback whales are capable of a more complex society than was once thought.

**Fig. 1.**
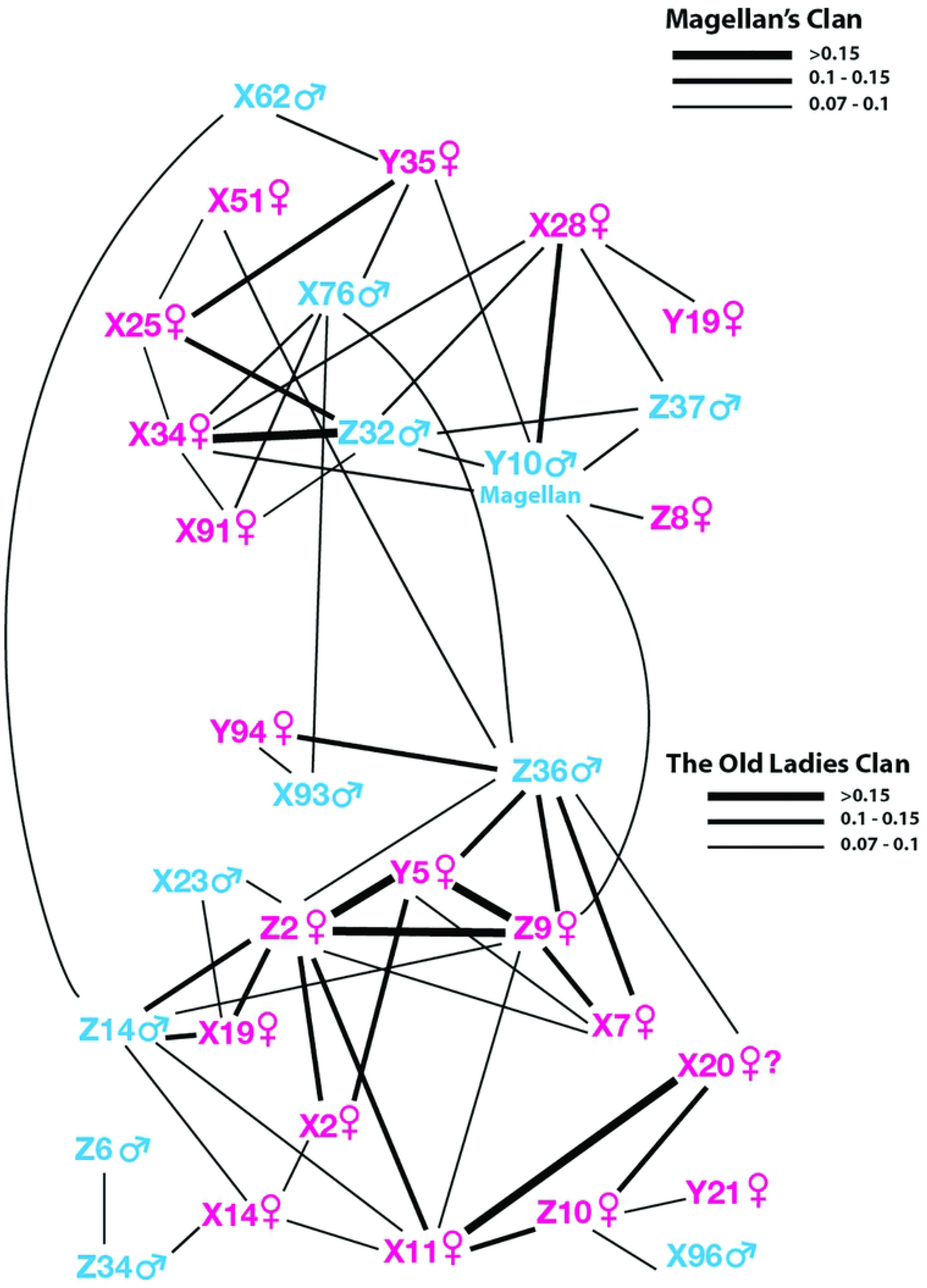
Prince William Sound in the northern apex of the Gulf of Alaska

## Methods

### Study area, surveys and effort

Observations were done with a scientific research permit under the Marine Mammal Protection Act (MMPA) and the Endangered Species Act (ESA) issued by NOAA’s National Marine Fisheries Service. (Amy Hapeman, NOAA Fisheries, Office of Protected Resources, U.S. Department of Commerce Office: 301-427-8419). All permit guidelines were adhered to at all times. Research was strictly observational with no biopsy or tagging, no handling, touching, or breaking the skin of animals. The Institutional Animal Care and Use Committee (IACUC) approval was not required.

The study area was comprised of 150 square miles of southwestern PWS including two large ocean entrances to the Gulf of Alaska. A centrally located research camp (permitted by the Chugach National Forest) was maintained on an island with expansive views (Fig. 2). Each year from May through October, surveys were performed using a small (7-9 meters) research vessel (RV). A “research day” consisted of at least 8 hours of survey time per RV. At times, multiple RV’s were used, therefore, the number of “research days” varied annually between 29 and 252 [16]. Whales were found by searching sometimes with the help of reported whale sightings. Weather limitations of winds of over Beaufort 4 or incessant rain halted searches. Vessel track lines and locations of whale encounters were drawn on data sheets. After 2003, GPS (Global Positioning System) was also used to document vessel track lines and latitude and longitude of each whale sighting. All animals were approached from behind at a distance of about 25 meters. As the target whale dove the pigmentation patterns on the underside of its tail were photographed with an SLR camera outfitted with a 200-400 mm lens. These photographs, termed “fluke shots” were used to identify individual whales [14].

**Fig 2.**
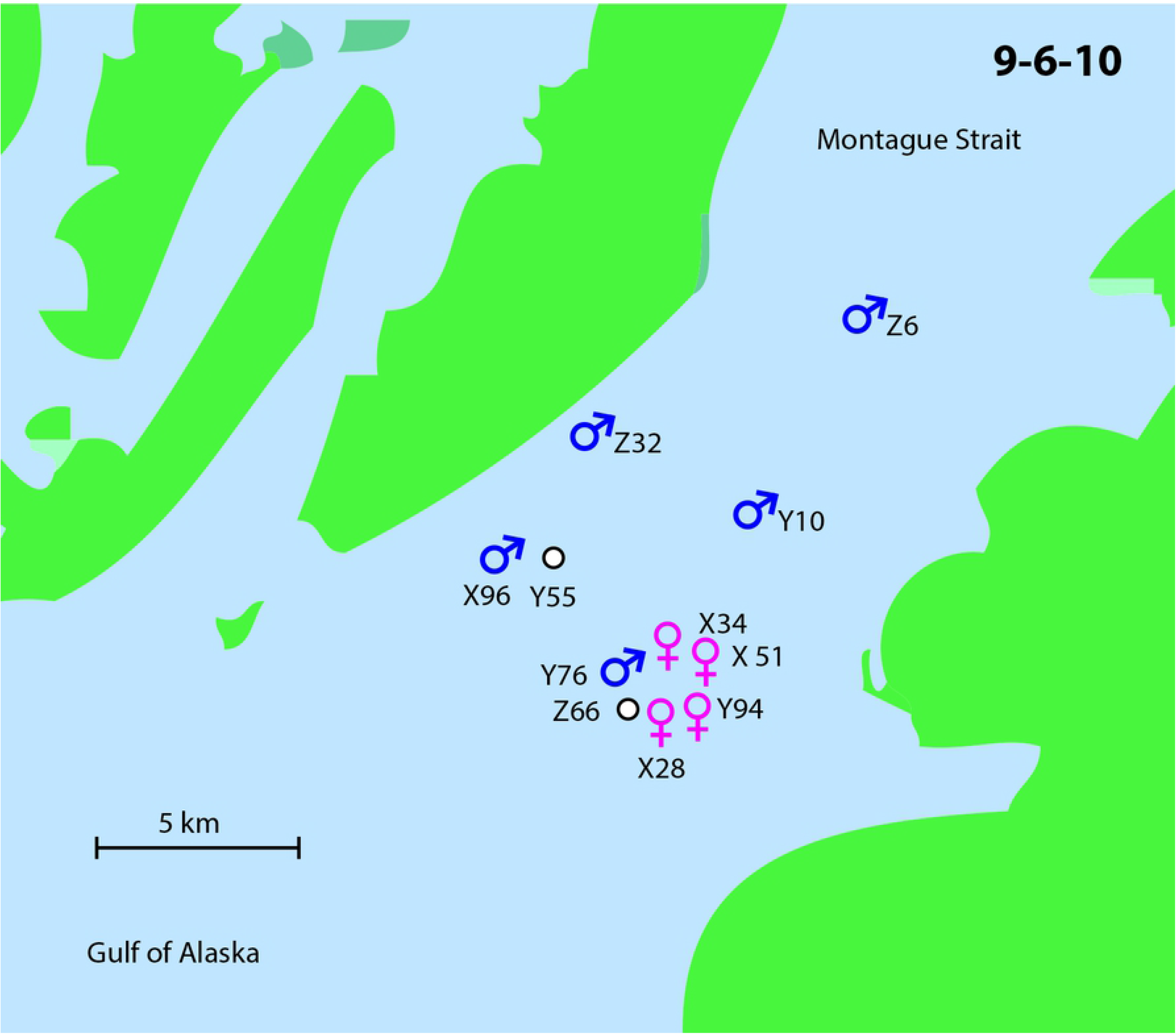
Study area with location of the field camp and 3 sub regions

### Sex determination

During the study period (1983-2009) there were 3,017 whale encounters involving 405 unique whales. Forty-six animals (26 females and 20 males) were sexed by DNA mitochondrial haplotype sampling [17]. Thirty-nine adult whales that were observed accompanied by a calf were assumed to be female. Seven whales, having at least an 8 consecutive year sighting history without a calf, were assumed to be males [18].

### Association analysis

Groups of humpback whales were often geographically isolated in small bays or passages. It was most common to find a few whales deep diving over the same food source with pairs often interchanging. All the individuals in such a group were termed “in association”. In larger bodies of water whales were also considered “in association” even if they were not geographically isolated but were in close proximity and displaying coordinated movement such as feeding, diving, resting, or traveling.

For consistency, the association analysis was based on humpback whale sightings from 3 geographic regions most reliably surveyed over the course of this study (Fig 2). Area 1 included Knight Island Passages and the bays immediately around the field camp. Area 2 included Montague Strait and Area 3 encompassed the ocean entrances. Sightings were grouped into 4 time periods: 1 (1983-1988), 2 (1989-1995), 3 (1996-2002), and 4 (2003-2009). This restricted the assessment of associations to time periods in which pairs of whales were present in the study area and enabled the investigation of changes in humpback whale associations across time. The fact that 1 time period had 6 instead of 7 years was statistically insignificant. An individual whale was considered to have a good sighting history within a given time period if it was observed a minimum of 7 times over 3 or more years within that period. Only individuals with good sighting histories for at least 2 consecutive time periods were included in the analysis. SOCPROG software (version 2.3) [19] [20] was used to calculate “simple ratio” association indices for all pairs of whales. This index ranges from 0 to 1.0 and reflects the proportion of all sightings of either whale in which both whales were observed associated with one another. All coefficients of .07 and higher were included in the analysis. A sociogram (Fig. 3) plotting the structure and strength of relationships was drawn.

**Fig. 3.**
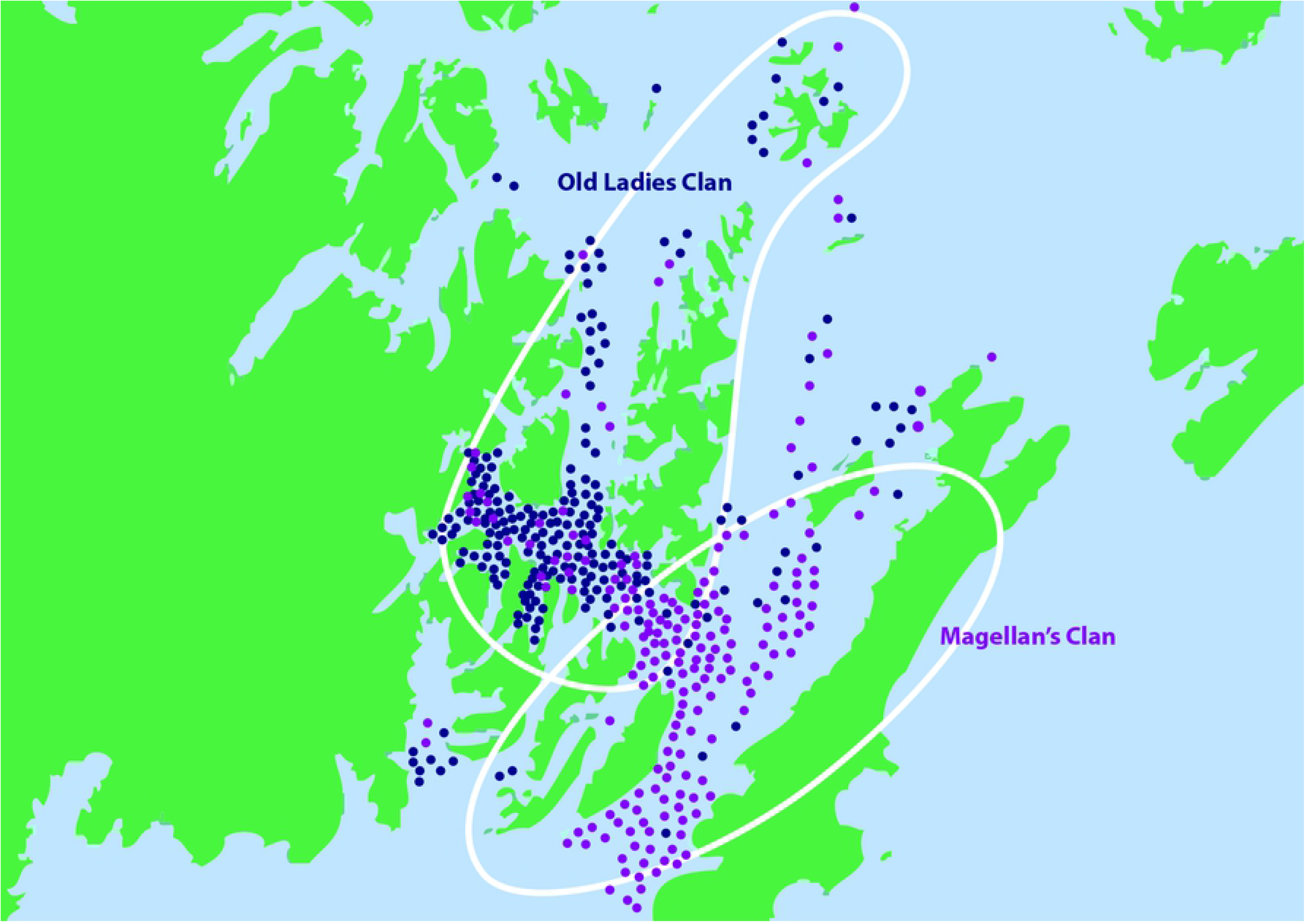
Sociogram of the two groups

### Migration matches

Fluke photographs were matched at the North Pacific Marine Mammal Laboratory and the SPLASH catalog [21] maintained by Cascadia Research Collective.

## Results

### Association analysis

Forty of the total 405 unique whales (9.88%) had long sighting histories, showing strong site fidelity. All associations of 0.07 were statistically significant, as were standard errors of association indices in the global data set (associations of individuals across all four time periods). The data therefore supported using a 0.07 threshold. Sixty strong and consistent associations were documented over 2 to 4 time periods between 32 whales. Two distinct groups were found and named the “Old Ladies Clan” and “Magellan’s Clan” (Table 2) (Fig. 3).

**Table 1.** Effort per year with days and nautical miles (NM)

| Year | Days | NM |
| --- | --- | --- |
| 1983 | 29 | 2305 |
| 1984 | 135 | 6124 |
| 1985 | 65 | 2404 |
| 1986 | 55 | 2527 |
| 1987 | 29 | 1111 |
| 1988 | 73 | 2331 |
| 1989 | 215 | 8738 |
| 1990 | 252 | 10585 |
| 1991 | 187 | 8451 |
| 1992 | 134 | 5665 |
| 1993 | 72 | 3019 |
| 1994 | 83 | 3413 |
| 1995 | 125 | 5976 |
| 1996 | 92 | 4158 |
| 1997 | 126 | 5722 |
| 1998 | 98 | 4451 |
| 1999 | 98 | 4514 |
| 2000 | 83 | 4001 |
| 2001 | 87 | 4272 |
| 2002 | 107 | 5226 |
| 2003 | 79 | 3722 |
| 2004 | 142 | 7785 |
| 2005 | 112 | 4686 |
| 2006 | 61 | 2696 |
| 2007 | 71 | 4013 |
| 2008 | 54 | 2128 |
| 2009 | 64 | 2640 |

**Table 2.**
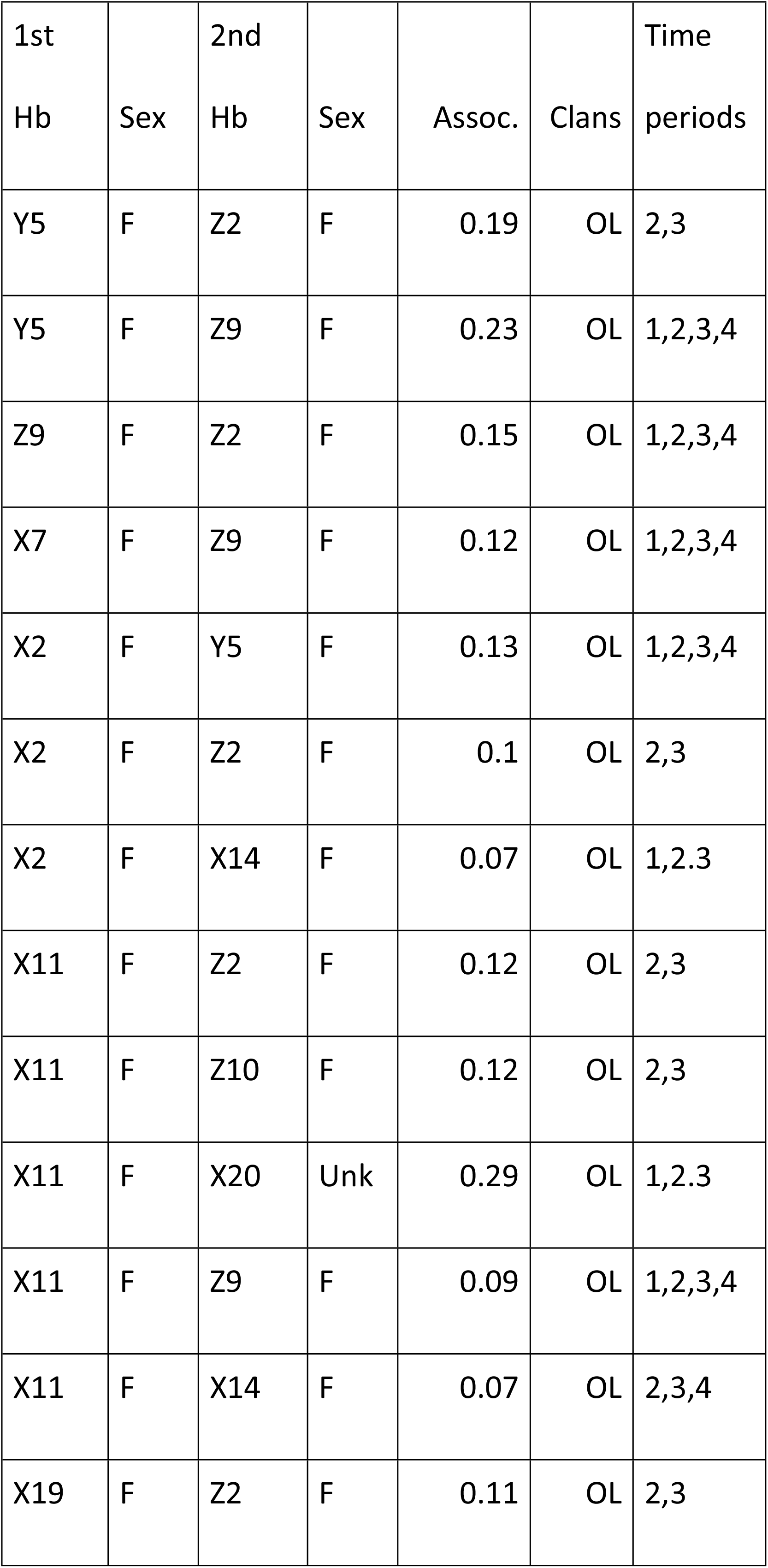
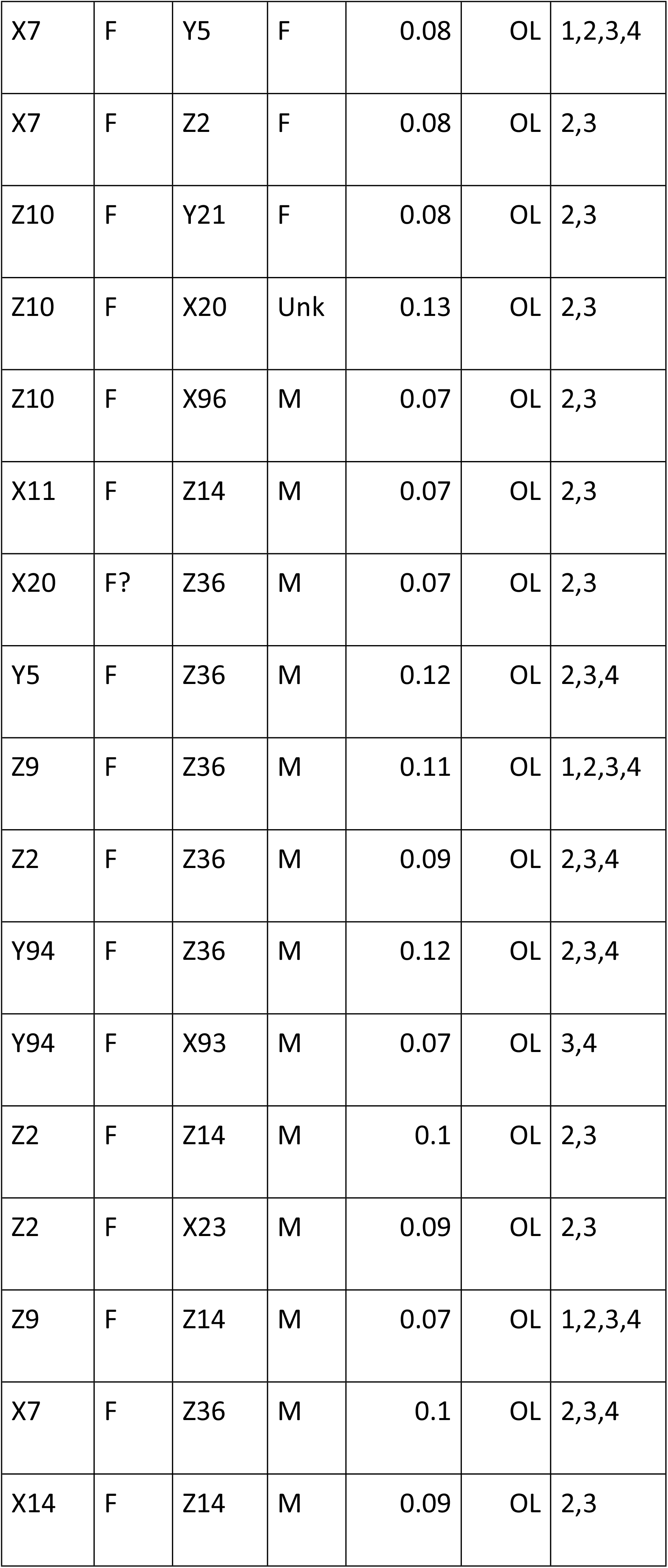
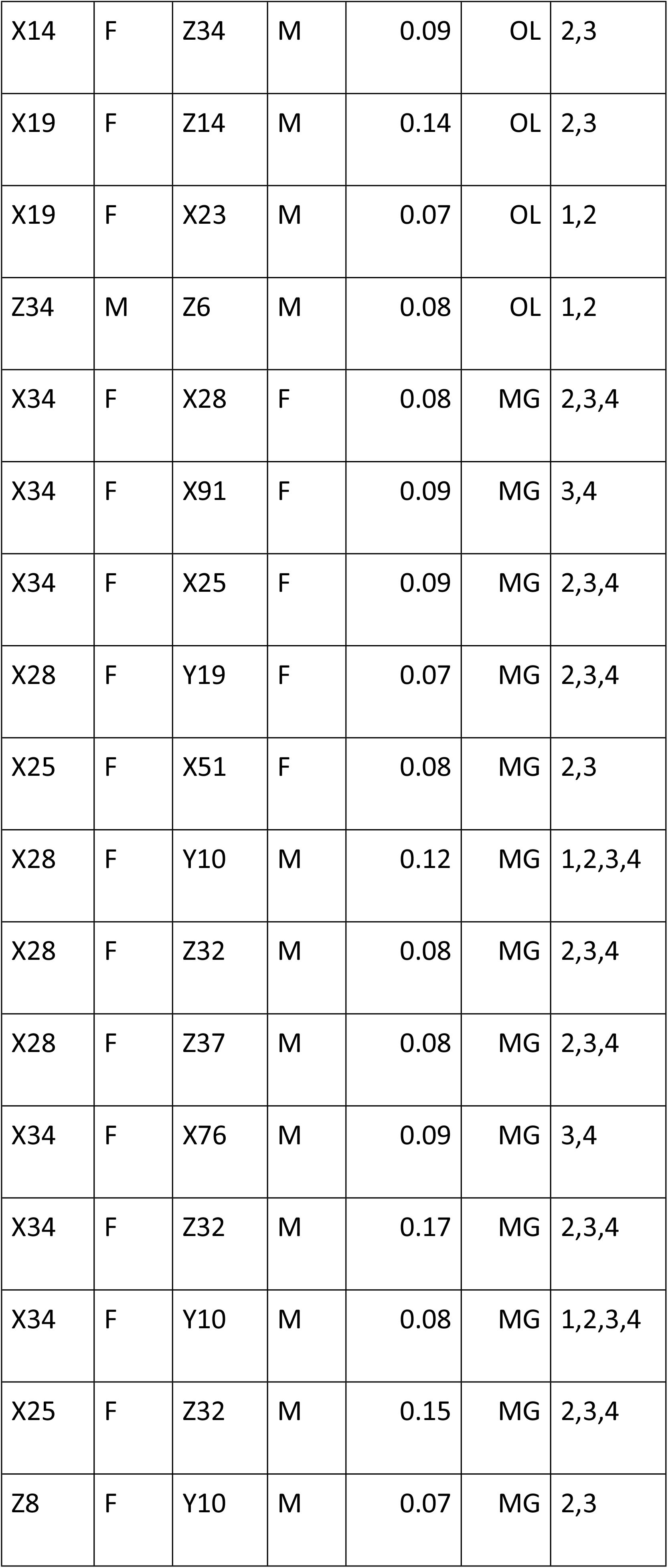
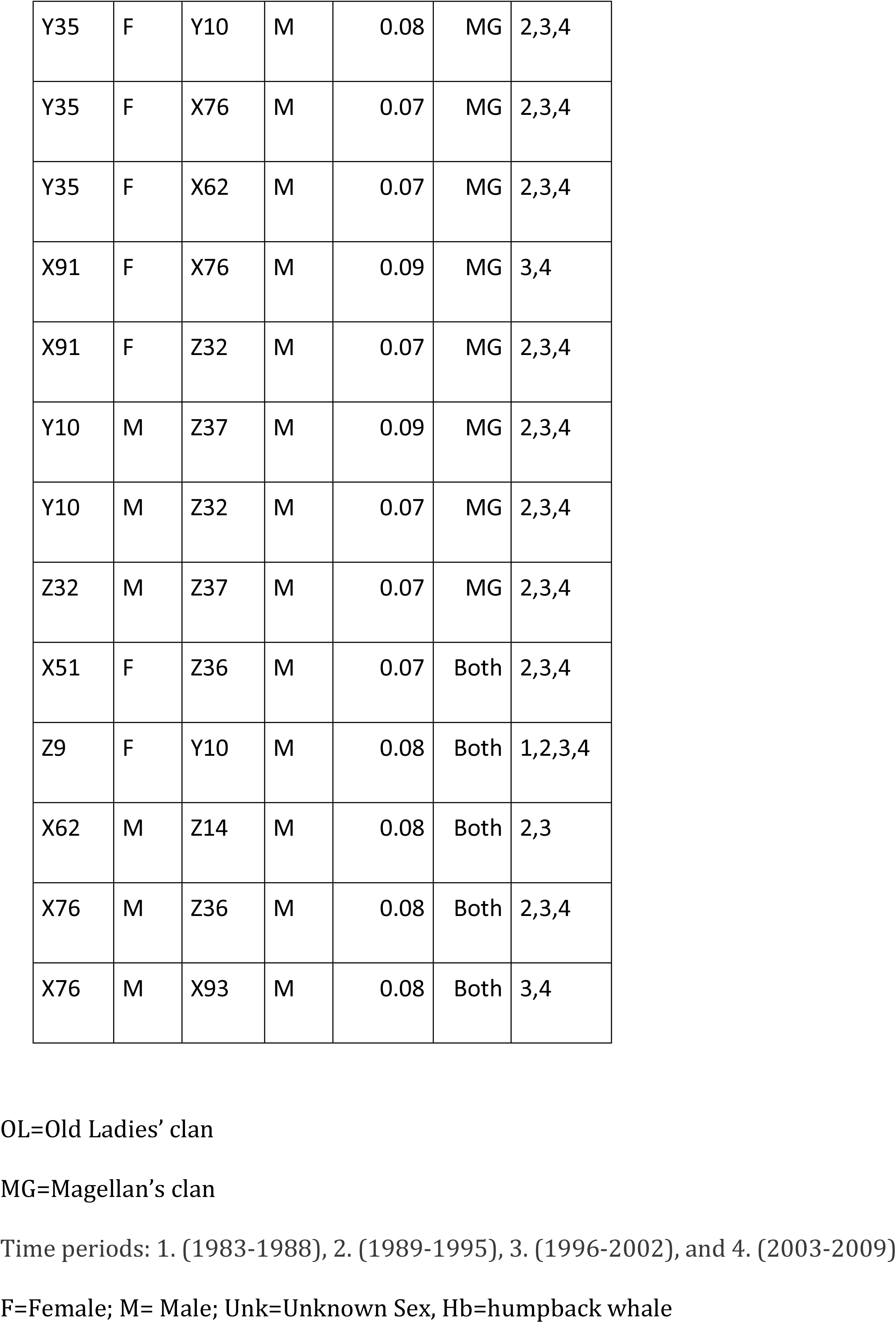
Association coefficients

### The Old Ladies Clan

The Old Ladies’ Clan consisted of 12 females and 7 males (0.08 to 0.29) over 2-4 time periods (13-27 years). Fourteen associations were female pairs (.1 to .29) and 13 were male/ female pairs (.07 to .14). One male-male association was found to have a lasting bond (with a coefficient of .08) over 3 time periods. Five older females stopped producing offspring during the study. The association coefficients were higher between females; however, even the oldest females had at least 1 associate male (Fig. 3).

### Magellan’s Clan

The second group of 13 whales, “Magellan’s Clan”, was named after a male that had a long sighting history in PWS and Baja California, Mexico [21][22]. This group had significant associations among 5 males and 8 females (.07 to .15). As in the other group, most of the females had associations with at least 1 male; however, the female-female associations in Magellan’s Clan were generally not as strong as those of the Old Ladies Clan. Five associations between females (.07 to .09) and 3 between males (.07 to .09) were significant (Table2) (Fig. 3).

There were 5 associations that crossed over between groups: 2 between a male and a female (.07 and .08) and 3 between 2 males (all .08) (Fig. 3). Six whales (2 males and 4 females) with very long sighting histories in the study area showed only weak ties to either clan and were called “Outliers”.

Sixty-five females were documented with 148 different calves of which 43 were successfully identified by fluke photographs in their first year with their mother. Only 9 of these young whales were re-sighted in PWS as adults [16]. Seven were offspring of the clan whales but were not observed associating with their maternal clan. Two (2) were calves of Thumbelina (Y2) (an “Outlier”). Only her son Brandon (X96), born in 1996, had strong ties with 1 female (Z10) in the Old Ladies Clan (Table 2) (Fig. 3).

Males were more often found alone (19% of 763 sightings) than females (12.77% of the 2,254 total sightings). Males of the clans were sometimes found alone at a distance from their affiliates (Fig. 4). These males eventually joined the others and often made a pass near the research vessel while it maneuvered among associate whales.

**Fig. 4.**
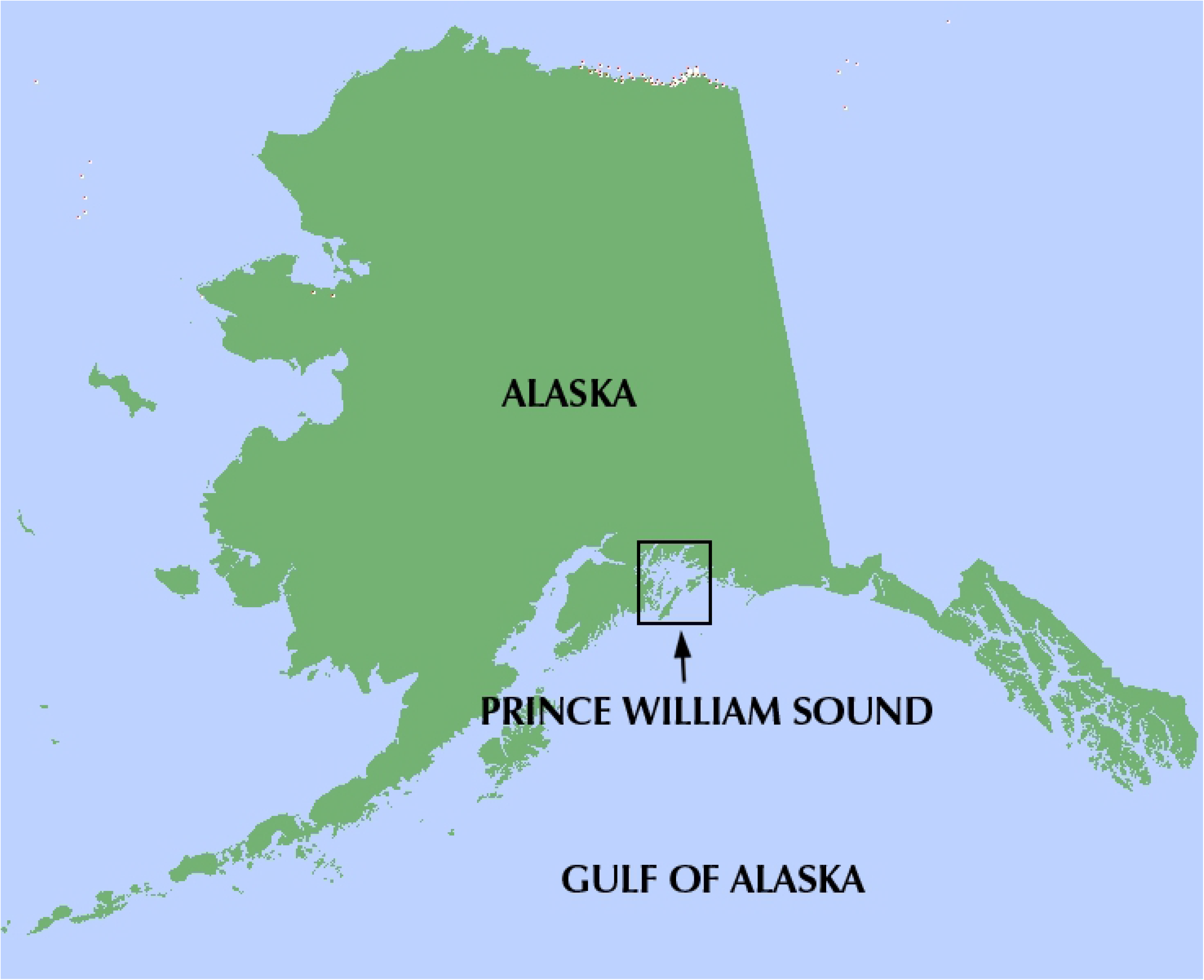
Example of spacing of clan males

### Ranges of the two clans

The Old Ladies Clan was mainly found in Knight Island Passage (Area 1). Magellan’s Clan was seen in Montague Strait and south to the Gulf of Alaska. (Area 2 and Area 3) and the lower part of Knight Island Passage where there was an overlap of clan ranges. Sightings of 2 males and 2 females from each clan were mapped to illustrate the ranges of sightings of the 2 clans (Fig. 5).

**Fig. 5.**
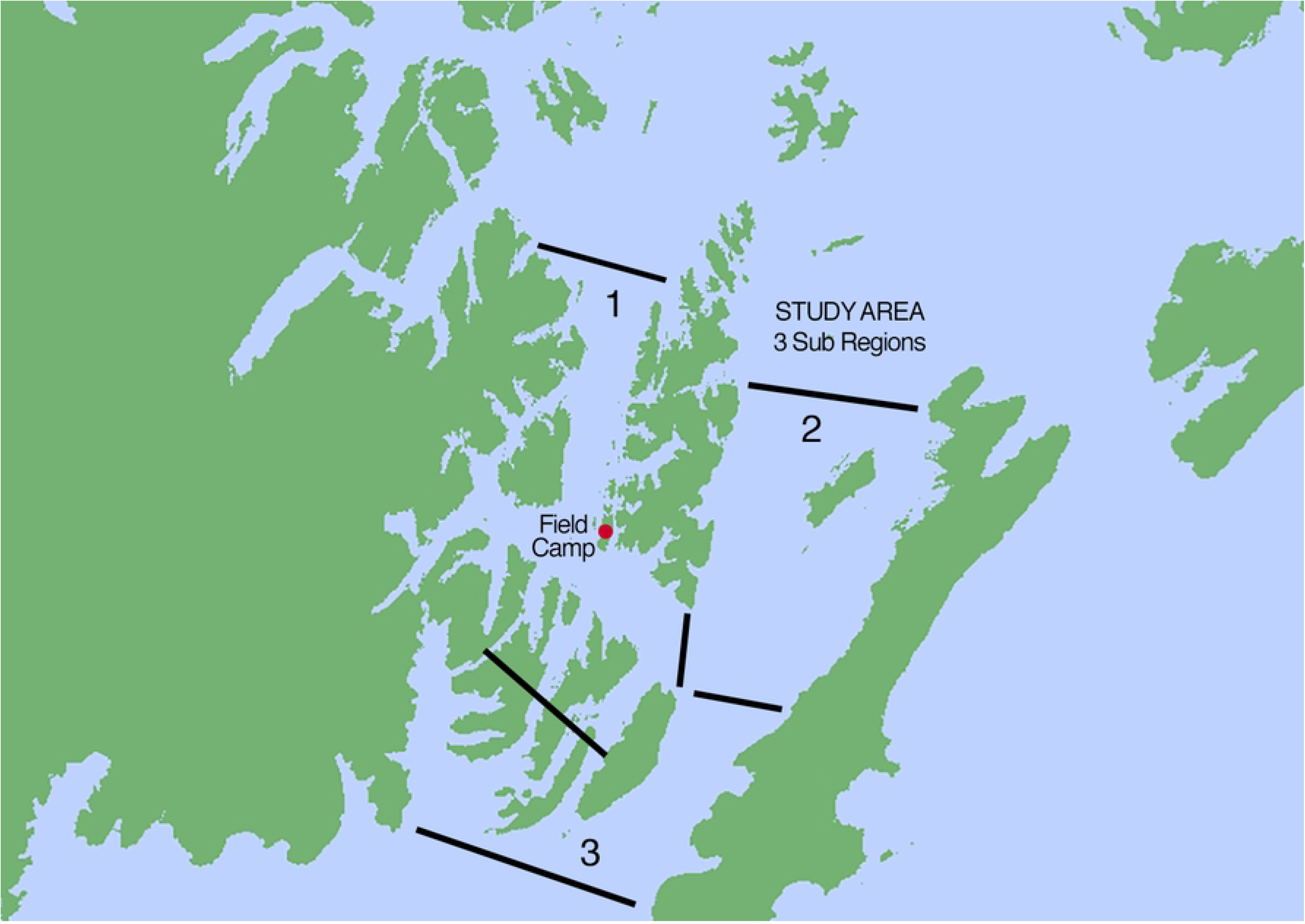
Ranges of clans

### Migration matches

Seven whales from Magellan’s Clan and 5 from the Old Ladies Clan have been photographed in Hawaii. Magellan (Y10) has been seen at least 3 times in Mexico Two whales from the Old Ladies Clan were documented swimming side by side off the coast of Maui on April 15, 2004 [21]. Female “Olgita” (Z10) had a newborn calf alongside and male “Skippy” (Z34) swam closely with the cow-calf pair acting as her “escort”.

## Discussion

The roles of male, female, old and young whales in humpback societies are not well understood. Their fluid fission-fusion [23–24] social behavior makes the endurance of long-term associations difficult to recognize and quantify. Our study in PWS represents an intimate view of a small population of humpback whales in their feeding grounds over a span of almost three decades. The two humpback communities described in PWS may represent local core groups of a much bigger stock with a wider range extending out into the North Gulf of Alaska. During the course of our study we observed some “new” whales peacefully feeding among the described “clan” whales while others caused obvious reactions when attempting to join. If the whales of these clans were part of a bigger social network this would explain the ability of some, but not all animals, to easily intermingle with these enduring groups.

Humpback whales must eat about 1,000 pounds of small schooling fish or krill (family Euphausiidae) per day [11][25]. In PWS, due to the depth of the fiords, most foraging whales are observed deep diving alone, in pairs, or small groups. Two distinct “clans” of individuals, each foraging in different parts of the study area, were recognized by the highest degrees of association. We suggest that strong enduring bonds amongst groups of individuals within larger populations of humpback whales may be important for the overall health, integrity, and survival of the species.

It is difficult to define an association in marine mammals because most of their behavior takes place underwater. Communication between humpback whales, through social sounds, is not well understood and may occur over many miles of ocean. Darling [26] suggests that whales within two kilometers of each other in both the breeding and feeding grounds could be associating. Whitehead [19] defines an associating group as a set of animals using a small area with suitable habitat that clearly separates the group. Mann [27] suggests defining “association” as including a behavioral condition “such as feeding” or “coordinated movement”. Often, an association between humpback whales has been defined as 2 or more whales swimming or diving synchronously side by side over a certain period of time [9][11][27]–[32]. PWS is comprised of many fiords and bays therefore groups of humpback whales were most often geographically isolated, as Whitehead describes. In this study coordinated movement was also used as a measure of association.

Males clearly had enduring associations with both males and females. It was once thought that male humpback whales were promiscuous, competing for females, with no long-term associations [1]–[5] [7] [29]. The idea that male-male associations in the breeding grounds are always competitive is now being questioned; males seem to be joining with other males displaying non-antagonistic behavior, possibly even cooperating to gain the ability to mate with a female [7][33]–[36]. This study shows associations between males enduring a number of years. Tyack [37] describes pairs of male dolphins forming long-lasting bonds and creating their own unique signature calls when they leave their mothers and form a new social group. The male humpback role may also have more importance in the health of a community than previously thought. In PWS nearly all the females had at least one affiliate male over many years. The strong enduring bonds between male humpback whales and the older females in this study suggest that non-producing females are still valued and may add integrity to their community. Strong female bonds may be beneficial not only for the individuals but also for the entire group. The bond between “Whitey” (Z2) and “Gwenivere” (Y5), increased throughout the 4 time periods (27 years). Their bond may have helped protect them from predation and may have increased their feeding success. Bonds amongst female humpbacks are found in the Gulf of Maine [9], the Gulf of St. Lawrence [32], and Southeast Alaska [18]. Strong female bonds are recognized in many animal societies such as primates [38], elephants [39], lionesses [40], and killer whales [41]. Long-lasting relationships among females in many animal societies are the basic unit of social life: lowering stress levels, ensuring safety from predators for individuals and their offspring, and increasing foraging success.

We suspect that some members of these groups migrate together to ensure strength and safety. Male “Skippy” (Z34) was discovered acting as an escort to female “Olgita” (Z10) off Maui, Hawaii. Both had been identified as members of the Old Ladies Clan in PWS. This was probably not coincidental and suggests these group bonds are more binding than previously thought. Further matching of fluke photographs taken in other regions could show how significant these long-term whale associations are between the feeding and breeding areas. Baker *et al.* [42] distinguishes a “central” North Pacific stock of humpback whales migrating to Hawaii for breeding and Alaska for feeding. Within this central stock are there smaller breeding stocks? Could there be a distinct PWS breeding stock? Based on sequences of DNA haplotypes (a group of alleles of different genes on a single chromosome) Witteven *et al.* [17] found PWS whales to be unique among whales in other Alaskan feeding ground sub regions (Kodiak, Southeast Alaska, and PWS offshore). Was “Skippy” protecting “Olgita” and her calf because he was protecting his unique gene pool? His community? Or even his offspring? We suspect that within these small social clans the male-female associations may have an importance that strengthens the group, with males protecting females young and old. We found that males sometimes stayed on the outer edges, or even a distance of a mile or two away from a cluster of their clan whales, however, at some point during the observation they would pass through the group. These males may have been protecting the clan by scouting for predators or intruders.

It was first thought that humpback whale offspring return to their maternal feeding grounds [12][43]–[49]. However, many have found that reproductive female humpback whales do not tend to associate closely with their own mature offspring [11] [30] [44]–[49]. In this study we found a few, but not all, grown offspring became regulars to PWS and these young adults did not show long-term bonds with their mothers. They also did not have strong associations with whales of the two social clans described in this study. Because many humpback whales move in and out of the Sound, the PWS calves may become part of other social groups feeding in other parts of the North Gulf coast. It may take many years for a young whale to establish enduring adult associations.

This study is different than most in that the same Principal Investigator performed observations on a small population of whales in isolated sheltered waters for nearly three decades. Two enduring clans were recognized. The authors suggest that similar social clans exist throughout humpback whale populations. Core groups have been described in Southeast Alaska [2] [8] [45] [30], however, because of the fission-fusion nature of humpback whale associations and the larger size of other populations and study areas, the value of small long lasting social clans may be more difficult to identify.

## Conclusion

Humpback whale social structure is complex and sustained over decades. They associate in distinct enduring clans with higher degrees of associations than previously thought. The strength and endurance of long-term whale associations, we believe, are building blocks of community: insuring securety, greater foraging success, and ultimately their survival.

## Acknowledgements

Thank you Ken Norris, who believed in us and sent us on our way in 1980 to Prince William Sound, Alaska to camp on a desolate beach and study and move amongst the whales with our small inflatable boat. And thanks to our parents for letting us go! Since then many people have helped through the years: volunteering on the research boat and at the field camp, reporting whales from their vessels, delivering fuel and food, and donating financial and moral support.

Craig Matkin, Eva Saulitas, Kathy Heise and Lance Barrett-Lennard from The North Gulf Oceanic Society (NGOS) provided fluke photographs and meticulous sighting data. Gerry Sanger (photographer and tour guide) also contributed numerous whale photographs and sighting information. Dean Kildaw did the analysis (SOCPROG) on our decades of data, and he, Dr. Deborah Boege-Tobin and Erin McMullan helped with the editing of this paper. Thank you all so very much.

